# *Phytophthora* methylomes modulated by expanded 6mA methyltransferases are associated with adaptive genome regions

**DOI:** 10.1101/217646

**Authors:** Han Chen, Haidong Shu, Liyuan Wang, Fan Zhang, Xi Li, Sylvans Ochieng Ochola, Fei Mao, Hongyu Ma, Wenwu Ye, Tingting Gu, Lubing Jiang, Yufeng Wu, Yuanchao Wang, Sophien Kamoun, Suomeng Dong

## Abstract

Filamentous plant pathogen genomes often display a bipartite architecture with gene sparse, repeat-rich compartments serving as a cradle for adaptive evolution. However, the extent to which this “two-speed” genome architecture is associated with genome-wide epigenetic modifications is unknown. Here, we show that the oomycete plant pathogens *Phytophthora infestans* and *Phytophthora sojae* possess functional adenine N6- methylation (6mA) methyltransferases that modulate patterns of 6mA marks across the genome. In contrast, 5-methylcytosine (5mC) could not be detected in the two *Phytophthora* species. Methylated DNA IP Sequencing (MeDIP-seq) of each species revealed that 6mA is depleted around the transcriptional starting sites (TSS) and is associated with low expressed genes, particularly transposable elements. Remarkably, genes occupying the gene-sparse regions have higher levels of 6mA compared to the remainder of both genomes, possibly implicating the methylome in adaptive evolution of *Phytophthora*. Among three putative adenine methyltransferases, DAMT1 and DAMT3 displayed robust enzymatic activities. Surprisingly, single knockouts of each of the 6mA methyltransferases in *P. sojae* significantly reduced *in vivo* 6mA levels, indicating that the three enzymes are not fully redundant. MeDIP-seq of the *damt3* mutant revealed uneven patterns of 6mA methylation across genes, suggesting that PsDAMT3 may have a preference for gene body methylation after the TSS. Our findings provide evidence that 6mA modification is an epigenetic mark of *Phytophthora* genomes and that complex patterns of 6mA methylation by the expanded 6mA methyltransferases may be associated with adaptive evolution in these important plant pathogens.

## Introduction

DNA methylation, one of the fundamental epigenetic marks, participates in many biological processes in both eukaryotes and prokaryotes^1-3^. The most studied form of DNA methylation is 5-methylcytosine (5mC), which is a prevalent DNA modification in mammals and plants^4^. The 5mC modification plays a role in many processes, such as transposon silencing, regulation of gene expression and epigenetic memory maintenance^5^. The amount of 5mC present in DNA varies across organisms and is barely detectable or absent in many species, such as the nematode (*Caenorhabditis elegans*), the fruit fly (*Drosophila melanogaster*) and brewers yeast (*Saccharomyces cerevisiae*)^6^. Comparatively, the N6-methyladenine (6mA) modification is extensively distributed in prokaryotic genomes. A prominent function of 6mA is in discriminating between host DNA and invading DNA, thus contributing to prokaryote immunity against phages and other invading genetic elements^7^. Besides, 6mA is also involved in DNA replication, repair, virulence, and gene regulation^8-11^.

In contrast to prokaryotes, the occurrence and biological functions of 6mA methylation in eukaryotic organisms remain largely uncharacterized. There is increasing evidence that 6mA is present in eukaryotes, including mammals, nematodes, algae, fruit flies, frogs, and fungi^12-16^. Genome-wide 6mA distribution patterns can be identified by several robust methods such as methylated DNA immunoprecipitation sequencing (MeDIP-seq)^14,15^, 6mA-sensitive restriction enzyme digestion coupled with high-throughput sequencing^17^, and single molecule real time sequencing (SMRT sequencing)^12,13^. The 6mA pattern appears to be dynamic during development; for instance, the early embryonic stage of *Drosophila* has relatively higher 6mA levels compared to later stages^14,18^. Furthermore, the genomic localization of 6mA significantly differs among organisms^13-15^. The 6mA modification is widely and evenly distributed in the *Caenorhabditis elegans* genome. By contrast, 6mA is enriched around transcription start sites (TSS) in early-diverging fungi and *Chlamydomonas*, and is enriched in transpoable elements in *Drosophila*. The localization patterns associate with 6mA biological functions. For example, in *Chlamydomonas* and fungi, 6mA is enriched around the TSSs of actively expressed genes, suggesting that 6mA may be an active mark for gene expression^12, 15^, while 6mA appears to suppress transcription on the X chromosome in mouse embryonic stem cell^16^

Like many other epigenetic marks, 6mA can be reversibly modulated by enzymes such as methyltransferase and demethylase^13,14^. It is known that DAM and M.MunI are classical bacterial 6mA methyltransferases^19^. In eukaryotic cells, enzymes from the MT-A70 protein family that evolved from M.MunI^20^, are considered 6mA methyltransferases. Overexpression of the MT-A70 homolog DAMT-1 from *C. elegans* in insect cells elevated the 6mA level, whereas knockdown of *damt-1* resulted in a decrease in the amount of 6mA, suggesting that DAMT-1 is a potential 6mA methyltransferase in nematodes^13^. However, methyltransferase-like protein 3 (METTL3) and METTL14 of the MT-A70 family catalyse 6mA on mammalian mRNA but weakly on DNA^21^. The Alkylation repair homologs (AlkB) protein family is involved in DNA damage repair, and could catalyse demethylation of both methylated DNA and RNA^13,16,22,23^. MT-A70 and AlkB homologs are prevalent in many organismsm and most of them are not functionally characterized. However, it is possible that other RNA and DNA demethylase and methyltransferase proteins could have evolved to regulate 6mA DNA in eukaryotic species.

The Oomycetes are a group of eukaryotic organisms that include a variety of pathogens that infect plants and animals^24^. A notorious example is *Phytophthora infestans,* the causal agent of potato late blight disease which sparked the Irish famine, resulting in starvation and migration of millions of people in the 1840s^25^. An additional example is *Phytophthora sojae*, a soybean root pathogen that currently threatens global soybean production. These two species are model organisms among oomycetes^26^. The genomes of these *Phytophthora* display a bipartite architecture, with gene-sparse and repeat-rich regions (GSR) and gene-dense regions (GDR) ^25^. The GSR compartments are associated with accelerated gene evolution, serving as a cradle for adaptive evolution^27-29^. However, the biological roles of DNA modifications and their associations with adaptive genome evolution remain unknown. In this study, we demonstrate that 6mA, rather than 5mC, is the major DNA methylation in these two *Phytophthora* species. We show that *P. infestans* and *P. sojae* genomes encode expanded numbers of 6mA methyltransferases (DAMT). Two of the three DAMTs have methyltransferase activity, and the 6mA methylation landscapes are described at the genome-wide level using methylated DNA immuno precipitation sequencing (MeDIP-seq). Although the majority of the methylation sites localized in the intergenic regions, 6mA also prefers to accumulate around TSS regions in a bimodal distribution pattern and may function as a repressive mark of gene expression. The GSR genes show higher a methylation level than the GDR genes. Consistently, most 6mA sites accumulate in repetitive sequences, such as DNA elements and long terminal repeat (LTR) elements. Furthermore, individual knockouts of each of the three *DAMT* genes results in a reduction of 6mA level *in vivo*. Moreover, comparative analysis of the MeDIP-seq data of the mutants suggests that the *DAMTs* may have functional specificity in targeting particular genomic regions.

## Results

To determine whether *Phytophthora* species can accomplish the 5mC modification, we performed a hidden Markov models based sequence similarity search for 5mC methyltransferase homologs in the *P. infestans* and *P. sojae* genomes^30,31^. No predicted gene or homologous sequence corresponding to a 5mC methyltransferase was discovered **(Supplementary Table 1**). To test directly for the presence of 5mC, we analyzed hydrolyzed genomic DNA (gDNA) samples from *P. infestans* and *P. sojae* by high-performance liquid chromatography (HPLC) and Ultra Performance Liquid Chromatography Electrospray Ionization - Mass Spectrum (UPLC-ESI-MS/MS). We did not detect 5mC in either species at the parts per billion (PPB) level (10^-9^g/mL) (**Supplementary Fig. 1a, b**). Furthermore, the endonuclease McrBC that specifically cleaves DNA containing 5mC did not digest *Phytophthora* gDNA, similarly to gDNA of *Drosophila melanogaster*, which is known to not carry 5mC (**Supplementary Fig. 1c**). Therefore, none of the methods we employed could detect 5mC in either *P. infestans* or *P. sojae* DNA.

Although we did not identify genes encoding for 5mC methyltransferases in *P. infestans* and *P. sojae*, we did identify homologs of 6mA methyltransferases and demethylases in the *Phytophthora* genomes. Initially, we discovered a potential MT-A70 homolog in the *P. sojae* but not *P. infestans* genome. However, closer examination of the putative *P. sojae* MT-A70 gene indicated that it is a pseudogene with a premature stop codon. We found that N6-adenineMlase domain-containing (*DAMT*) proteins are present in all the examined oomycete species, including *Phytophthora* species, *Albugo* species, *Hyaloperonospora arabidopsidis*, *Pythium ultimum* and *Saprolegnia parasitica* (**Fig. 1a**). The *Phytophthora* and *Saprolegnia* genomes each encode three predicted *DAMT* genes, whereas the other species have only one gene. Phylogenetic analyses of the oomycete DAMTs uncovered two distinct gene clades, namely DAMT1/2 and DAMT3 (**Supplementary Fig. 2a**). In contrast to *DAMT1* and *DAMT2*, *DAMT3* is conserved in all the examined oomycete genomes except *H. arabidopsidis* (**Supplementary Fig. 2a, Supplementary Table 2**). *DAMT3* is located in a genomic region with a high degree of synteny (**Supplementary Fig. 2b**), suggesting that it is probably the ancestral gene. *DAMT* gene expansion in *Phytophthora* species therefore appears to be due to the emergence of the *DAMT1/2* genes. A closer examination of the catalytic motif responsible for binding the methyl group from S-adenosyl-L-methionine ^13,20,32^ indicates that DAMT1 and DAMT3 proteins have functional motifs consisting of the amino acid sequences DPPY and DPPF, respectively. However, this motif was naturally mutated into EPPH in the DAMT2 proteins. A search in the *P. infestans* and *P. sojae* online RNA-seq databases revealed that *DAMTs* are expressed in all the examined growth stages^33,34^ (**Supplementary Fig. 2d, e**). In summary, bioinformatics analyses indicate that *Phytophthora* species may possess the enzymatic machinery for 6mA DNA methylation.

**Figure 1.**
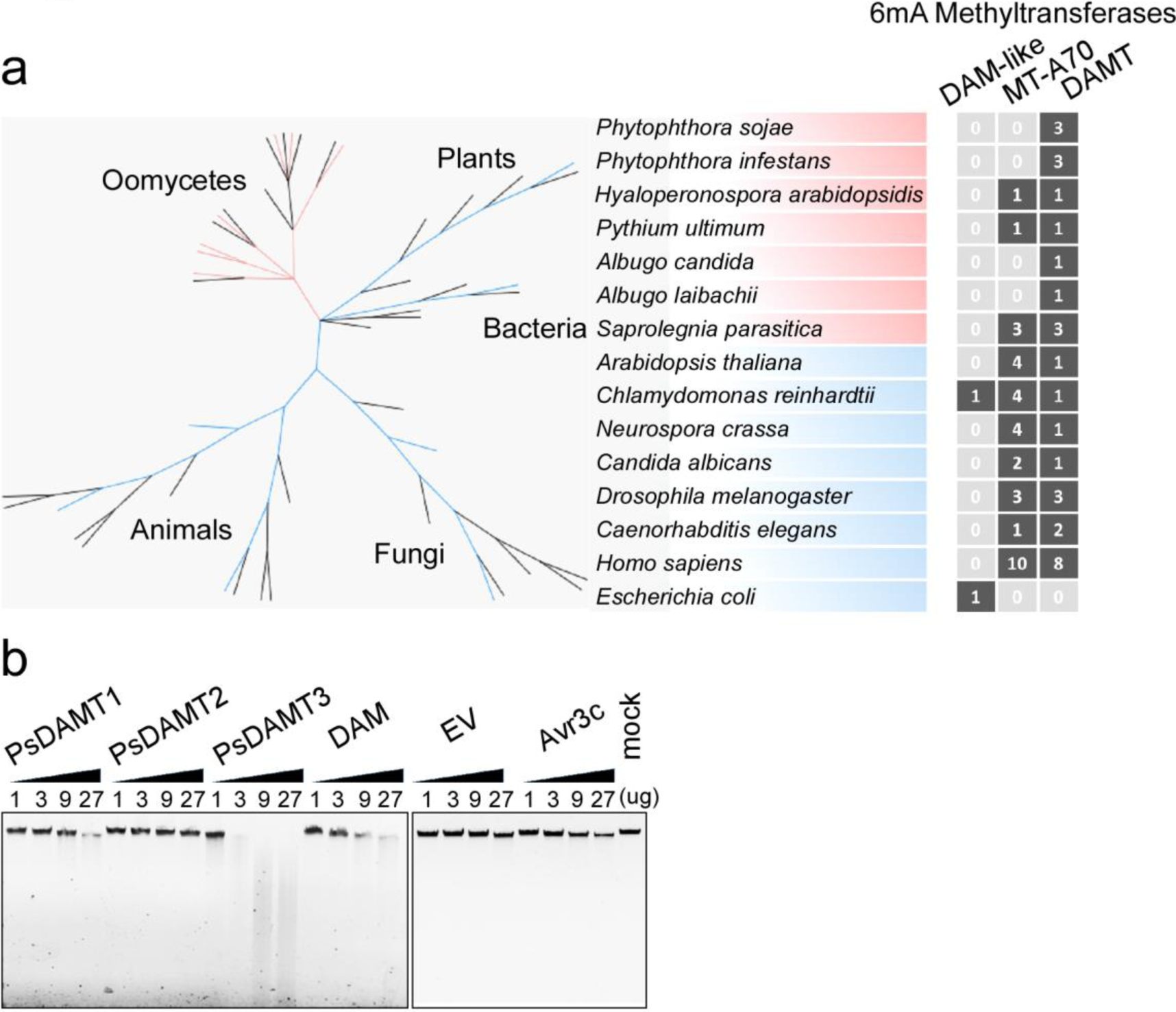
*Phytophthora* genomes encode functional 6mA Methyltransferases. **(a)** N6-adenineMlase domain-containing (*DAMT*) gene expansion in *Phytophthora* species. Two families of methyltransferase genes (*DAM-like*, *MT-A70*) and the *DAMT* family genes from seven oomycete organisms and eight model organisms are shown in a simplified phylogenetic tree. Color codes: oomycetes (red), other model organisms (blue), presence of gene homolog (black) and absence of gene homolog (grey). The number of homologous genes in each organism is also labelled. **(b)** *In vitro* DpnI-dependent DNA methylation assay suggests that *Phytophthora* DAMTs have methyltransferase activity. Recombinant proteins PsDAMT1, PsDAMT2, PsDAMT3 together with bacteria 6mA DNA methylase (DAM) were produced in *E. coli*. EV (empty vector) and Avr3c (a *Phytophthora* secretion protein) were used as controls. The recombinant protein gradient ranged from 1 µg to 27 µg in each reaction. The experiments were carried out by triplicates with similar results.

To verify the enzymatic activity of these putative methyltransferases, we measured the *in vitro* methyltransferase activity of three *P. sojae* recombinant DAMT proteins. The recombinant proteins, together with 6mA-free lambda DNA and substrate S-adenosyl-L-methionine, were incubated together in an *in vitro* enzymatic assay^35^. These assays revealed that lambda DNA is smeared by treatment with the restriction enzyme DpnI, which recognizes the 6mA methylated GATC site, in the presence of recombinant PsDAMT1, PsDAMT3, or the bacterial 6mA methyltransferase DAM (**Fig. 1b**). Notably, PsDAMT3 was the most active methyltransferase *in vitro*. We did not detect any activity for PsDAMT2 in this assay, even after increasing PsDAMT2 concentration (**Fig. 1b**). We also performed a complementary methylation assay in the 6mA deficient *E. coli* strain HST04.

In this assay, *E. coli* gDNA from DH5α and *DAM* complemented HST04 transformants were digested by DpnI as expected. The *E. coli* gDNA from *PsDAMT1* and *PsDAMT3* transformants could also be digested by DpnI, whereas those from *PsDAMT2* and the transformants of the catalytically dead mutants (*PsDAMT1^APPA^*, *PsDAMT3^APPA^*) could not be digested (**Supplementary Fig. 3a**). Overall, these data indicate that the *P. sojae* DAMT1 and DAMT3 proteins possess methyltransferase activity in a DPPY(F) motif-dependent manner.

To test for the presence of 6mA in *Phytophthora*, we used UPLC-ESI-MS/MS to analyze gDNA samples from *P. sojae* and *P. infestans*. A peak matching the retention time of standard 6mA was present in the test samples from these two *Phythophthora* species (**Fig. 2a**). Moreover, the same base fragment was detected in the two samples by MS/MS of the 266.12 (mass/charge ratio), which also matched the standard 6mA (**Fig. 2b**). Thus, the 6mA DNA base modification is present in the gDNA samples. Furthermore, we estimated the abundance of 6mA in *P. sojae* and *P. infestans* to be 400 and 500 parts per million (PPM), respectively, as determined by UPLC-ESI-MS/MS (**Fig. 2c**, **Supplementary Table 2**). The 6mA level in these *Phytophthora* species is approximately 60-fold higher than in *Homo sapiens* and *Mus musculus*, but is lower than a few early-diverging fungal species like *Hesseltinella vesiculosa* and *Piromyces finnis*^12,36^. To further test for the presence of 6mA in *Phytophthora* gDNA, we used commercially available antibodies that specifically recognize the 6mA modification; immune blot signals were robustly detected in gDNA samples of *P. infestans* and *P. sojae.* (**Fig. 2d**). Collectively, our results show that 6mA is a naturally occurring DNA modification in *Phytophthora* genomes.

**Figure 2.**
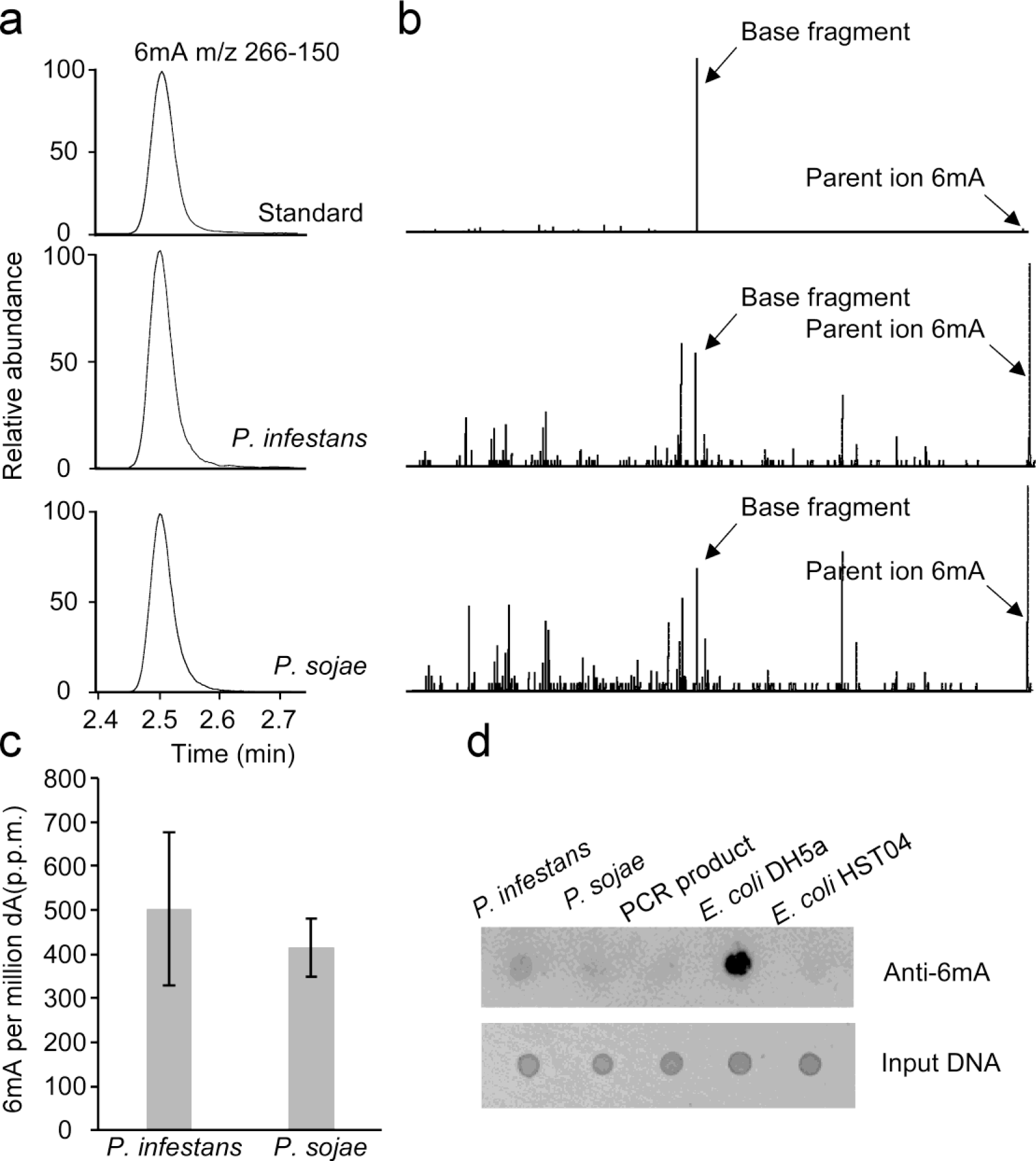
6mA occurs in *Phytophthora* genomic DNA. **(a)** *Phytophthora* mycelium 6mA were detected by UPLC. The selective multiple reaction monitoring (MRM) transitions for 6mA were setting as m/z 266-150. The retention time of standard 6mA was present in *Phytophthora* gDNA samples. **(b)** *Phytophthora* 6mA are detected by UPLC-ESI-MS/MS. The parent ion 6mA (m/z near 266.12) and base fragment (m/z near 150.07) highlighted by arrows from samples also matches standard 6mA. **(c)** Quantification of 6mA levels in *Phytophthora* samples. The 6mA concentrations are listed as 6mA per million dA. **(d)** The presence of 6mA in *Phytophthora* gDNA was verified by dot blot assay using a specific 6mA antibody. Input DNA was quantified by ethidium bromide-dyed agarose gels. Every dot contained 100 ng DNA. The experiments were independently carried out in triplicate.

We performed methylated DNA immuno precipitation-sequencing (MeDIP-seq) to obtain a genome-wide insight into the *Phytophthora* 6mA methylome. The MeDIP-seq experiments on gDNA samples from mycelium growth stages included two biological replicates for each of the two *Phytophthora* species. After assembling sequencing data and seeking 6mA-enriched regions, we mapped 6mA peaks (6mA-enriched regions) at a genome-wide level with FDR < 0.01 by SICER^37^. A total of 12,611 overlapping methylation peaks were captured from the two *P. infestans* biological replicates. A total of 3,031 overlapping peaks were called from two *P. sojae* replicates (**Supplementary Fig. 4a**). Genome-wide 6mA methylation profiling data revealed that 86% and 55% of the 6mA peaks were located in the intergenic regions in *P. infestans* and *P. sojae*, respectively (**Supplementary Fig. 4b**). The higher proportion of 6mA intergenic localization in *P. infestans* results from the larger overall fraction of intergenic gDNA in the expanded 240 Mbp genome of this species compared to *P. sojae*. In *P. sojae*, 25% of the 6mA peaks mark gene bodies, whereas 15% and 5% of the methylations occupy positions upstream and downstream of gene bodies, respectively. Comparatively, in *P. infestans*, these figures correspond to 8%, 4%, and 2% (**Supplementary Fig. 4b**). Overall, our analyses revealed 1,805 and 1,343 genes with 6mA marks in *P. infestans* and *P. sojae*, respectively.

Profiling of 6mA distribution in methylated genes revealed that 6mA peaks tend to flank the transcriptional start site (TSS) with a clear depletion near the TSS itself (**Fig. 3a-c**), resembling the bimodal distribution pattern of 6mA detected in other organisms, such as *Chlamydomonas*^15^. This bimodal distribution pattern can be verified by heatmap analyses when we plot relative 6mA levels from all the methylated and non-methylated genes (**Fig. 3c**). We illustrate normalized 6mA MeDIP-seq reads mapped onto loci from *P. sojae Ps_155563*, *Ps_128235* and *P. infestans PITG_02506*, *PITG_02507*, *PITG_15808* as typical examples of 6mA localization patterns (**Supplementary Fig. 5a, b**). To gain further insight into the characteristics of 6mA methylated genes, we conducted a gene ontology (GO) enrichment analysis of methylated genes in both species. Results from the GO analysis suggest that methylated genes are associated with functional categories, such as chromatin binding, enzyme regulator, and hydrolase (**Supplementary Fig. 5c**).

**Figure 3.**
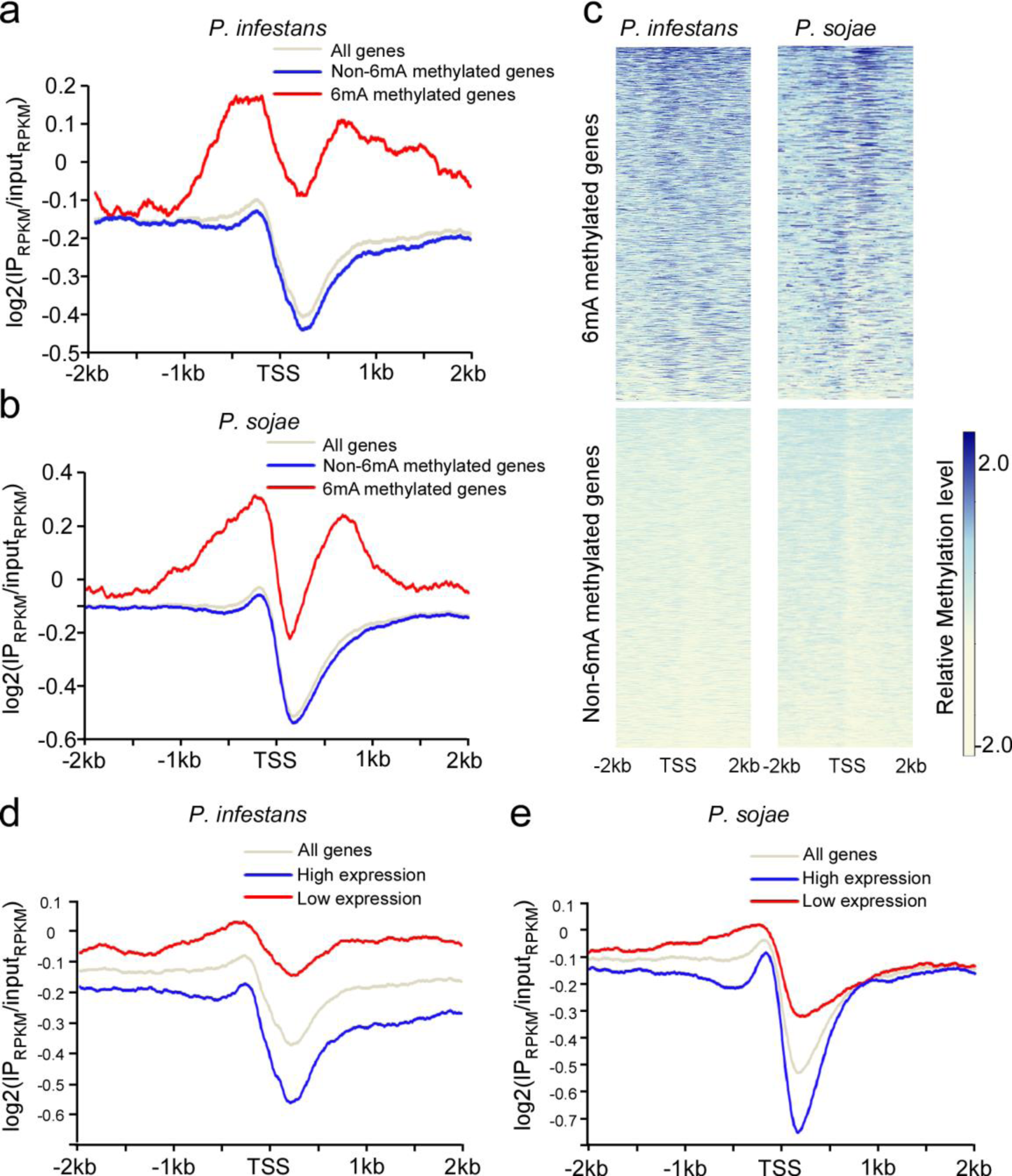
Bimodal distribution pattern of 6mA around TSS in *Phytophthora*. The distribution of 6mA peaks around TSS was profiled by MeDIP-seq. The 6mA occupancy along TSS from −2kb to 2kb is shown. 6mA peaks were enriched around TSS with a bimodal distribution and a local depletion after TSS in *P. infestans* and *P. sojae*. **(a)** 6mA occupancy in *P. infestans* methylated genes. All *P. infestans* genes are divided into 6mA methylated genes (n=1805) and non-6mA methylated genes (n=16374). **(b)** 6mA occupancy in *P. sojae* methylated genes. All *P. sojae* genes are divided into 6mA methylated genes (n=1343) and non-6mA methylated genes (n=17853). **(c)** Heatmap analyses of 6mA signal from individual genes verified the bimodal distribution pattern in both *Phytophthora* species. The relative methylation signal is represented using gradient colors. **(d)** The 6mA level is negatively correlated with gene expression in *P. infestans*. All genes are divided into two groups: high expression (FPKM>5, n=9927) and low expression (FPKM<5, n=8252). **(e)** The 6mA level is negatively correlated with gene expression in *P. sojae*. All genes are divided into two groups: high expression (FPKM>5, n=9450) and low expression (FPKM<5, n=9746).

Although it is debatable how well a GO analysis can inform questions of biological function, there is increasing evidence that 6mA is an important epigenetic mark for the regulation of gene expression^12,15^. In particular, the bimodal localization of the 6mA signal around the TSS prompted us to investigate the relationship between 6mA modification and gene expression. We compared the 6mA gene methylation data with RNA-seq gene expression data and examined the average 6mA level of highly expressed genes (FPKM>5) and lowly expressed genes (FPKM<5) in *P. infestans* and *P. sojae*. Lowly expressed genes are more likely to be associated with 6mA as this group of genes tends to have more abundant 6mA levels; in contrast, highly expressed genes tend to have lower 6mA levels (**Fig. 3d, e**). To further validate these observations, we examined the gene expression levels of methylated and non-methylated genes in both species. We found that methylated genes have significantly lower gene expression compared to non-methylated genes in both species (**Supplementary Fig. 6a, b**). Thus, the data suggests that 6mA negatively correlates with gene expression levels in the two *Phytophthora* species.

It is well established that genomes of *Phytophthora* species have experienced repeat-driven expansions and are, therefore, rich in repetitive sequences^25-28^. Thus, we examined the association between 6mA peaks and major types of transposable elements (TEs). A total of 37% (*P. infestans*) and 15% (*P. sojae*) of the 6mA peaks locate to long terminal repeat (LTR) elements (class I TEs), whereas 8% (*P. infestans*) and 10% (*P. sojae*) *of* the peaks fall within DNA elements (class II TEs), respectively (**Fig. 4a**). Statistical analyses indicate that 6mA peaks are enriched in TEs at a significant level (**Supplementary Fig. 7a**). Moreover, 6mA levels in TEs are higher than the average genomic level in both species (**Supplementary Fig. 7b**). We conclude that the 6mA methylome is preferentially associated with TEs in the two *Phytophthora* species.

**Figure 4.**
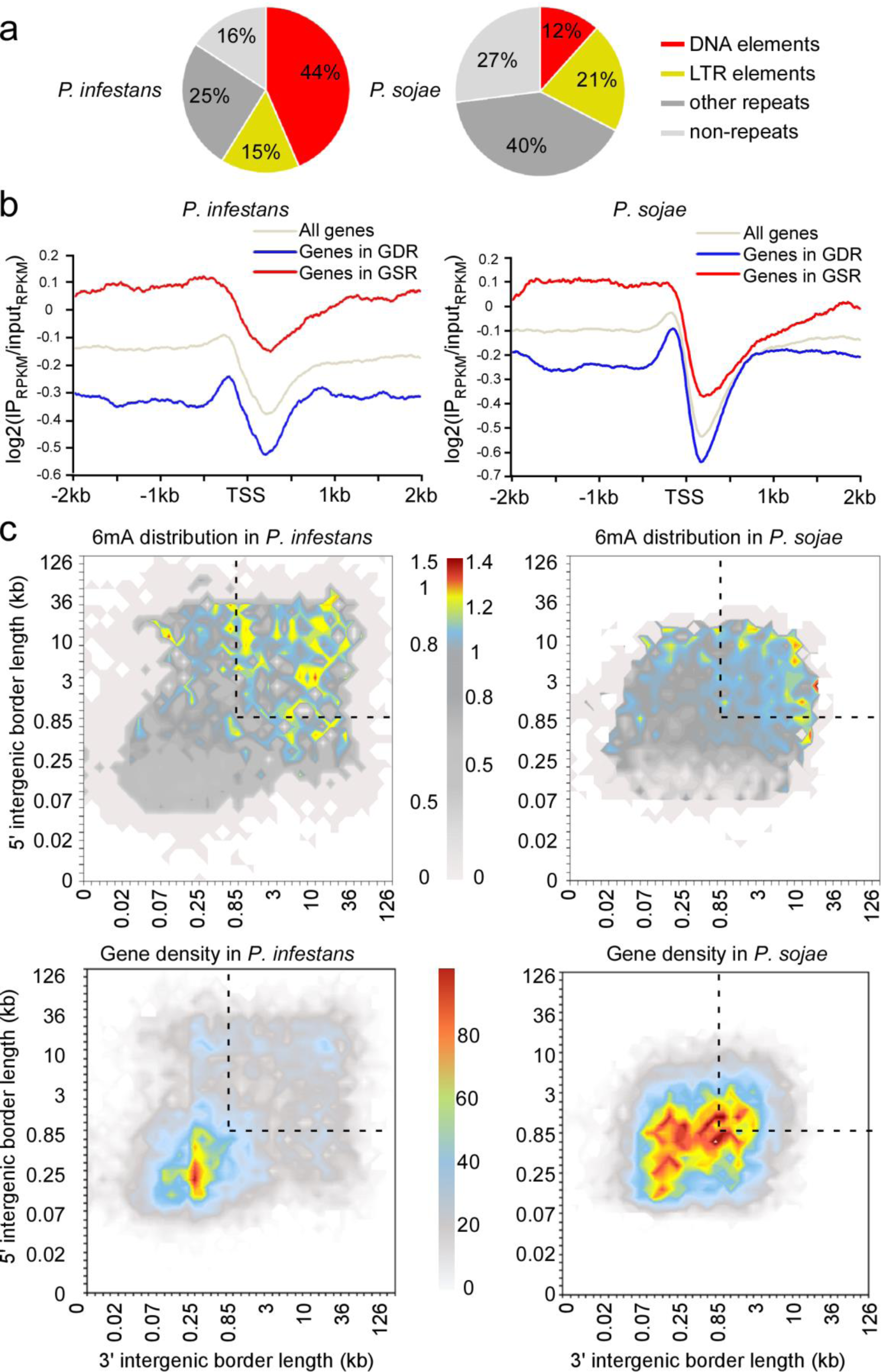
6mA marks *Phytophthora* repetitive elements and highlights gene sparse region. **(a)** Pie charts illustrate that 6mA peaks are predominantly distributed in repeat sequences in *P. infestans* and *P. sojae.* **(b)** Genes occupying the gene-sparse regions (GSR) display higher levels of 6mA than genes occupying the gene-dense region (GDR) in *P. infestans* and *P. sojae*. In *P. infestans*, GSR occupying genes (n=3920) and GDR occupying genes (n=6526) were calculated. In *P. sojae*, GSR occupying genes (n=3154) and GDR occupying genes (n=7240) were calculated. **(c)** Heatmap analyses reveal that 6mA accumulate in *P. infestans* and *P. sojae* GSR. All the genes in each genome were sorted into two dimensional bins on the basis of the lengths of flanking intergenic distances to neighbouring genes at their 5’ and 3’ ends. For the 6mA distribution heatmap, gradient color represents the average normalized 6mA value of genes in each bin. For the gene density distribution heatmap, gradient color represents the number of genes in each bin. The dotted line highlights GDR.

The genomes of *Phytophthora* species have a bipartite “two-speed” architecture with distinct gene dense regions (GDR) and gene sparse regions (GSR) ^25-28^. The dynamic GSR are enriched in rapidly evolving genes, such as virulence effectors, and these regions are thought to enable a faster rate of pathogen evolution^27-29^. To investigate the relationship between 6mA and genome architecture, we calculated the average 6mA levels for genes located in GDR and GSR. These analyses revealed that genes in the GSR tend to have higher 6mA level than GDR genes (**Fig. 4b, c**). Similarly, we plotted the 6mA methylation RPKM value of the region corresponding to the 500bp after the TSS according to local gene density (measured as length of 5’ and 3’ flanking intergenic regions) to generate the genome architecture heatmaps previously described^25,27^. The heatmaps revealed a clear association between the methylome and genome architecture that genes with higher 6mA levels being enriched in the GSR and reduced in GDR (**Fig. 4d, Supplementary Fig. 8a**). These observations are consistent with our previous finding that 6mA preferentially accumulate in repetitive and TE-rich regions, which fill the intergenic regions in the GSR of *Phytophthora* genomes. Interestingly, further MeDIP-seq analyses demonstrated that secretome genes, including RxLR effector genes, which are important in *Phytophthora*-host interactions and are primarily localized in the GSR, have significantly higher 6mA levels than core orthologous genes (**Supplementary Fig. 8b, c**). We conclude that the 6mA methylome is preferentially associated with both the genes and intergenic regions that form the gene-sparse compartments of *Phytophthora* genomes.

To further investigate the function of DAMTs in *Phytophthora*, we individually knocked out *DAMT* genes in the *P. sojae* strain P6497 using CRISPR/Cas9 gene editing methodology. We designed two sgRNAs matching two sites in each of the *DAMT* genes and harvested at least three independent knockout transformants for each gene (**Supplementary Table 3, Supplementary Figure 9**). We selected homozygous mutants *psdamt1*-T21 (-139bp), *psdamt2*-T52 (-1bp), and *psdamt3*-T9 (-374bp) as representative strains for further analyses. To examine the 6mA level in the *PsDAMTs* mutants, we applied UPLC-ESI-MS/MS to quantify 6mA abundance. 6mA levels in *psdamt1*-T21 and *psdamt2*-T52 were dramatically reduced to only 8.1% and 5.8% of the wild-type strain, whereas *psdamt3*-T9 was reduced to 11.0% (**Fig. 5a, Supplementary Table 4**). Although our biochemistry assays showed that PsDAMTs have different levels of enzymatic activities, these experiments show that each of the three *PsDAMT* genes significantly contribute to 6mA modification *in vivo*. Remarkably, the three enzymes do not appear to be fully redundant as each of the single knock-out mutants had a reduced methylation level.

**Figure 5.**
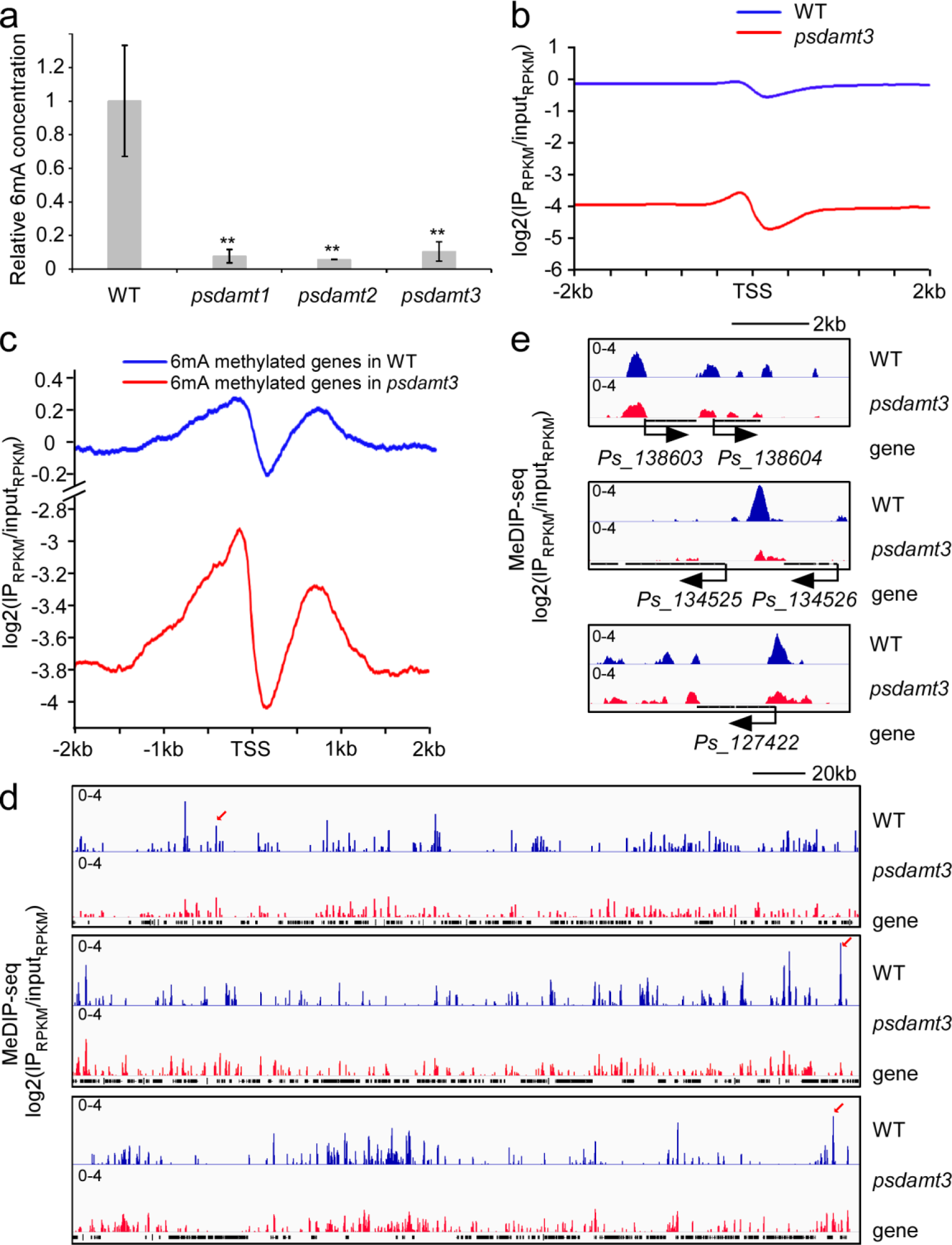
6mA level and landscape are altered in the *Phytophthora* DAMT3 mutant. **(a)** The 6mA levels of DAMT mutants were quantified by UPLC-ESI-MS/MS. WT is *P. sojae* strain P6497. *psdamt1* (T21), *psdamt2* (T52) and *psdamt3* (T9) are representative lines from each of the *DAMT* gene knockout mutants generated by CRISPR/Cas9. ** represents significant differences (P<0.01, Students’ *t* test) **(b)** The average 6mA level of all the genes (n=19196) is reduced in the *P. sojae psdamt3* mutant. **(c)** The average 6mA level of methylated genes (n=1343) is reduced in the *psdamt3* mutant. **(d)** Snapshot of 6mA deposition in the *psdamt3* mutant demonstrates that 6mA are widely but unevenly reduced in a representative genomic segment. **(e)** Zoom in snapshot of 6mA deposition in *psdamt3* at two gene loci as illustrated in (d).

Given that *DAMT3* encodes a functional methyltransferase that is conserved among all examined oomycete species, we examined the methylome in the *psdamt3* T9 mutant in more detail using MeDIP-seq. We observed a significant reduction in 6mA levels around TSS (**Fig. 5b**). 6mA signals were also weaker in *psdamt3* than wild-type at LTR elements and DNA elements regions (**Supplementary Fig. 10a**). Also, we noted a similar reduction in 6mA levels for both GSR and GDR genes in the *psdamt3* mutant compared to wild-type (**Supplementary Fig. 10b**). We conclude that the *PsDAMT3-*regulated 6mA methylome is not specifically associated with the bipartite genome architecture. However, close examination of the bimodal methylation pattern around the TSS of 6mA methylated genes uncovered a greater loss in the second peak in the *psdamt3* mutant compared to the wild-type (**Fig. 5c**). This unexpected finding indicates that *DAMT* genes may have some degree of functional specialization, and that PsDAMT3 may have a preference for the methylation of gene bodies after the TSS. Representative genomic segments with typical changes in 6mA localization are illustrated for *psdamt3* and wild-type *P. sojae* (**Fig. 5d, e**). The MeDIP-seq data partially explains the significant reduction of total 6mA levels in the *psdamt3* mutant, but also illustrates the uneven reduction pattern around the TSS, suggesting that there are complex patterns of 6mA methylation by the expanded 6mA methyltransferases of *P. sojae*.

## Discussion

It recently became evident that 6mA is not only an important epigenetic mark in prokaryotes but is also a feature of eukaryotic genomes. Here, we demonstrate that 6mA methylation occurs in the oomycete plant pathogens *P. infestans* and *P. sojae*, and document the methylome of these species. Remarkably, the 6mA methylomes are preferentially associated with genes and transposable elements that form the gene-sparse compartments of *Phytophthora* genomes and have been implicated in adaptive evolution of these pathogens. We discovered that 6mA methyltransferases have expanded into three enzymes in *Phytophthora* which do not appear to be fully redundant given that each of the single knock-out mutants had a significantly reduced methylome. Based on mutant analyses, we noted that PsDAMT3 may have a preference for methylation of gene bodies after their TSS. Overall, the observed 6mA patterns around the TSS in the *PsDAMT3* mutant suggest complex patterns of 6mA methylation by the expanded 6mA methyltransferases of *P. sojae*.

Although 6mA appears to be prevalent in eukaryote genomes, most studies report low levels of abundance. Previous studies documented significant variation of 6mA abundance (6mA/A) ranging from 0.00019% to 2.8% among different eukaryotes^12^. Here, we determined the abundance of 6mA to be 0.05% and 0.04% in the mycelium stage of *P. infestans* and *P. sojae,* respectively. Beside modification abundance, genome distribution pattern is another way to value 6mA biological significance. Unlike reports from some other organisms, 6mA is not evenly distributed across *Phytophthora* genome from our research. We revealed an unexpected link between 6mA methylation and the two-speed genome architecture of *Phytophthora* genomes. We noted an enrichment of 6mA peaks in the intergenic regions, particularly in repeat sequences such as DNA and LTR transposable elements. Other recent studies have suggested that 6mA participates in the regulation of transposon expression in *Drosophila* and mammals^14,16^. It has been recognized that proliferation of transposable elements could drive the adaptive genome evolution of filamentous pathogens, such as plant pathogenic fungi and oomycetes. To control the activity and the spread of these repetitive sequences, transposable-element rich regions are normally condensed with a high level of DNA methylation^38,39^. The genomes of *Phytophthora* species have a greater proportion of repetitive sequences compared to other oomycete species that have been sequenced to date^25^. We speculate that DAMT gene expansion in *Phytophthora* species was likely a consequence of transposable-element activity and that the occurrence of 6mA in repeat elements might function to inhibit the activity and spread of transposable elements in *Phytophthora* species. Therefore, 6mA may play a role in regulating genome integrity and plasticity to the optimal levels necessary for rapid evolution.

Recent studies documented that 6mA is associated with active genes in several organisms, and a popular model is that 6mA may associate with DNA/nucleosome structure to alter gene transcriptional processes^12, 15^. In *Chlamydomonas*, 6mA shows a bimodal localization pattern around TSS and frequently modifies DNA linkers between adjacent nucleosomes around TSS^15^. Studies in *Xenopus laevis* and *Mus musculus* found a marked decrease in 6mA in the vicinity of TSS^40^. 6mA is also predominantly distributed around TSS in a few fungal species^12^. In *P. infestans* and *P. sojae*, we observed a bimodal distribution pattern of 6mA enriched regions flanking the TSS, but with a clear depletion at the TSS itself and immediately downstream. This pattern resembles 6mA methylation described for the green algae *Chlamydomonas^41^.* Our comparisons of transcriptome and methylome data suggest that 6mA is negatively correlated with gene expression in the two *Phytophthora* species. Indeed, 6mA depletion is primarily located upstream of TSS in *Chlamydomonas* whereas it mainly located downstream of TSS in the two *Phytophthora* species. This could account for the apparent different associations of 6mA with gene expression in oomycetes and green algae. Our results are more reminiscent of a recent report in mammallian systems that 6mA is a negative gene expression mark in mouse embryonic stem cells^16^. This is consistent with our hypothesis that 6mA inhibits transposon activity. It is possible that interplays between 6mA marks and other epigenetic modifications, transcriptional regulators and other factors that are enriched around TSS, work in concert to regulate gene expression. Further investigations are required to explore the roles of 6mA in the modulation of gene expression.

*Phytophthora* genomes are well known for their bipartite “two-speed” architecture with hundreds of gene sparse regions comprised of repeat sequences and virulence effector genes serving as a cradle for adaptive evolution^27-29^. *Phytophthora* genes in the GSR tend to have higher 6mA levels and are also enriched in plant-induced genes that are normally silenced *in vitro*^27,42^. Indeed, we observed higher 6mA levels in secretome genes and RxLR effector genes (**Supplementary Fig. 8b, c**). This data is also consistent with the observation that 6mA marks are associated with low gene expression levels and may therefore contribute to the global down-regulation of virulence effector genes during vegetative stages, as proposed for the fungal pathogen *Leptosphaeria maculans*^43^. Alternatively, 6mA marks may contribute to the stochastic gene silencing of effector genes, thus enabling the emergence of pathogen races that evade plant immunity. Indeed, effector gene silencing has been linked to rapid evolution in both *P. sojae* and *P. infestans*^44,45^. In addition, *Phytophthora* species tend to exhibit high levels of expression polymorphisms in genes located in the GSRs^46^. In summary, we hypothesize that 6mA is involved in virulence gene expression, thus shaping host adaptation and enhancing evolvability in the plant pathogen *Phytophthora*.

In this study, we identified genes predicted to encode 6mA methyltransferases and demethylases in *P. infestans* and *P. sojae*. We initially focused on studying the functionality of the predicted 6mA methyltransferases to provide evidence that *Phytophthora* species have the inherent capability to perform this DNA modification. Our present work shows that DAMT homologs are the major N6-adenine methyltransferases in *Phytophthora* species. Although MT-A70 homologous proteins function in performing 6mA methylation in *C. elegans*^13^, our findings suggest that MT-A70 type methylases do not participate in 6mA methylation in *P. infestans* or *P. sojae*. MT-A70 homologs are either missing or pseudogenized in the *Phytophthora* species we examined. Our results also indicate that DAMTs underwent gene expansion in *Phytophthora* species compared to related oomycete genera such as *Hyaloperonospera* and *Albugo*. Among the three putative methyltransferases we characterized, DAMT3 appears to be the ancestral gene and encodes the methylase with the highest *in vitro* activity. To our surprise, knockout of each of the *DAMT* genes in *P. sojae* resulted in a substantial and comparable reduction of 6mA abundance *in vivo*. Like the *psdamt3* mutant, 6mA abundance in the *psdamt1* and *psdamt2* mutants is reduced to a similar level, despite the differences we observed in the *in vitro* activity of each of the DAMT enzymes. The results suggest that all three DAMT genes are required for efficient 6mA methylation in *P. sojae*.

The observation that all three *Phytophthora DAMT* genes contribute to 6mA genome methylation is intriguing. Our MeDIP-seq data uncovered that altered 6mA signals from the *psdamt3* mutant are unevenly spread across the genome. This observation suggests that certain 6mA sites could be preferentially regulated by *DAMT1* or *DAMT2.* Meanwhile, it also indicates that 6mA gene body modifications after the TSS are preferentially produced by *DAMT3*. We propose that gene expansion may have led *Phytophthora* 6mA methyltransferases to specialize, and thus they may not be fully functionally redundant. Previous reports showed that RNA adenine methylase METTL3/METTL14 form a stable heterodimer core complex in human cells^21, 47^. It remains possible that *Phytophthora* DAMTs associate as a complex and function collaboratively *in vivo*. The mode of action of *Phytophthora* DAMTs in the methylation process and their roles in targeting particular genome compartments require further investigation.

The mechanisms underpinning epigenetic modifications in *Phytophthora* have remained poorly understood ever since the observations of internuclear spread of gene silencing by van West and colleagues almost 20 years ago^48^. DNA methylation inhibitor 5-azacytidine and histone deacetylase inhibitor trichostatin-A released the silencing state of the *inf1* elicitin gene in *P. infestans^48^*. Silencing Dicer-like, Argonaute, and histone deacetylase genes reversed the expression of sporulation gene *cdc14*^49^. More recently, naturally occurring gene silencing of an avirulence effector gene in *P. sojae* was associated with the appearance of small RNAs^44^. These data suggest that epigenetic regulation plays a role in virulence and development of *Phytophthora* species. Although DNA methylation is a common type of epigenetic modification in many organisms, the extent to which *Phytophthora* genomes are methylated has remained unclear. Van West and his colleagues failed to detect 5mC by bisulfite sequencing in an endogenous locus that is sensitive to DNA methylation inhibitor^48^. Our results not only clarify that 5mC is absent in *Phytophthora* species but also provide evidence that 6mA shapes the epigenetic landscape in this lineage of organisms. To our best knowledge this is also the first 6mA methylome report from stramenopile or heterokonts organisms. This work provides a starting point to further explore 6mA epigenetic regulation in oomycete organisms, with important implications for plant pathology and management of plant diseases. Our results together with emerging studies in other organisms suggest that 6mA fulfills distinct and perhaps differing roles across the spectrum of eukaryotic organisms.

## Methods

### Phytophthora and plant cultivation

*P. sojae* reference strain P6497 was routinely cultured on solid 10% V8 agar medium at 25 ⁰ C in the dark. Non-sporulating hyphae were cultured at 25 ⁰ C in the dark using 10% V8 liquid medium for 3 days. *P. infestans* T30-4 strain was routinely cultured on the solid RSA/V8 medium at 18 ⁰ C in the dark. Non-sporulating hyphae were cultured at 18 ⁰ C in the dark in 10% V8 medium for 6-7 days. Hyphae were collected and immediately frozen using liquid nitrogen. Soybean cultivar Hefeng47 and Williams were used to provide etiolated hypocotyl after growing at 25 ⁰ C (16h) and 22 ⁰ C (8h) for 4 days in the dark.

### Data sampling

For homologous protein search, we selected 23 sequenced species, including 15 oomycete species and 8 model organisms as shown in Fig.1a. Their genome sequences were downloaded from EnsemblGenomes (http://ensemblgenomes.org/) and Joint Genome Institute (http://genome.jgi.doe.gov/). N6-adenineMlase (PF10237), MT-A70 (PF05063), DAM (PF05869), DNA_Methylase (PF00145), and MethyltransfD12 (PF02086) from the PFAM database were used to BLAST search homologous enzymes with an E-value cut-off 10^-5 29,30^.

### Dot blot assay

Genomic DNA of *P. sojae* and *P. infestans* were extracted using TIANGEN DNAsecure Plant kit. Different amounts of gDNA were denatured at 95 ⁰ C for 5 mins and chilled in ice for 10 mins. DNA were spotted on HybondTM-N+ membranes. The membrane was allowed to dry at 37 ⁰ C for 20 mins and then crosslinked using HL-2000 HybriLinker for 5 mins. The membrane was blocked in 5% milk PBST for 1h at room temperature, and then incubated with 6mA antibody (sysy202003) in 5% milk PBST overnight at 4 ⁰ C. After 3x 10 min washes with PBST, DNA and membrane were incubated with secondary antibody (ab6721) for 30 mins at room temperature. After 3x 10 min washes with PBST, the membrane was treated with Pierce ECL Western Blotting Substrate (Prod#32106) and detected by Tanon-5200Mutil. 100 ng input DNA of every samples were loaded on 1% agarose gels, followed by air drying for 5 mins and photographed using Clinx GenoSens.

### HPLC analysis for 5mC

The HPLC separation was performed on a Zorbax SB-C18 column (2.1 mm x 150 mm, 5 mm, Agilent) with a flow rate of 0.8 mL/min at 30 ⁰ C. Methanol (with 0.1% Formic Acid, v/v, solvent A) and 10 mM potassium phosphate monobasic in water (with 0.1% Formic Acid, v/v, solvent B) were employed as mobile phase. A gradient of 3 min 90% B with a flow rate of 0.8 mL/min, 1 min 90% B with a flow rate of 0.8-0.2 mL/min, 11 min 90% B with a flow rate of 0.2 mL/min, 3 min 90% B with a flow rate of 0.2-1.2 mL/min, 10 min 90% B with a flow rate of 1.2 mL/min, and 2 min 90% B with a flow rate of 1.2-0.2 mL/min was used.

### UPLC-ESI-MS/MS analysis for 5mC and 6mA

Analysis of the DNA samples was performed on UPLC-ESI-MS/MS system consisting of a Waters Xevo TQ-S micro mass spectrometer (Waters, Milford, MA, USA) with an electrospray ionization source (ESI) and an Acquity UPLC-I-Class^TM^ System (Waters, Milford, MA, USA). Data acquisition and processing were performed using Masslynx software (version 4.1, Waters, Manchester, UK). The UPLC separation was performed on a reversed-phase column (BEH C18, 2.1 mm×50 mm, 1.7 μm; Waters) with a flow rate of 0.2 mL/min at 35 ⁰ C. FA in water (0.1%, v/v, solvent A) and FA in methanol (0.1%, v/v, solvent B) were employed as mobile phase. A gradient of 5 min 5% B, 10 min 5-30% B, 5 min 30-50% B, 3 min 50-5% B, and 17 min 5% B was used. The mass spectrometry detection was performed under positive electrospray ionization mode.

A HPLC-ESI-MS/MS system, consisting of an electrospray-time-of-flight mass spectrometry (Triple TOF 5600+, AB Sciex) and a liquid chromatography (LC-20ADXR HPLC, Shimadzu), was also used for 5mC detection. Data acquisition and processing were performed using PeakView version 2.0 (AB Sciex). The HPLC separation was performed on a reversed-phase column (C18, 2.1 mm×100 mm, 2.6 μm; Kinetex) with a flow rate of 0.2 mL/min at 40 ⁰ C. FA in water (0.1%, v/v, solvent A) and FA in methanol (0.1%, v/v, solvent B) were employed as mobile phase. A gradient of 15 min 20-90% B, 3 min 90% B, 0.1 min 90-20% B, and 1.9 min 20% B was used. The mass spectrometry detection was performed under ESI positive mode with a DuoSpray dual-ion source.

### DpnI-dependent methylation assay

DpnI-dependent methylation assay was performed as previously described^34^. The reaction contained 20 mM Tris-HCl, pH 8.0, 50 mM NaCl, 7 mM 2-mercaptoethanol, 1 mM EDTA, 0.1 mg/ml bovine serum albumin (BSA), 1 µg N6 -methyladenine-free lambda DNA, purified recombinant proteins (1-27 µg), and 50 µM unlabeled AdoMet. The reaction system were incubated at 37 ⁰ C for 1 hour, and then 65 ⁰ C for 15 min to stop the reaction. The methylated DNA was digested by 5U DpnI at 37 ⁰ C for 1 hour. Digestion was stopped by heat inactivation by incubating at 80 ⁰ C for 20 mins. 1% agarose gel electrophoresis was used to check digestion. PsDAMT1, PsDAMT2, PsDAMT3, DAM, PsAvr3c were cloned into pET32a-c (+). The recombinant plasmids were transformed into *E. coli* HST04 strain (*dam-, dcm-*). Bacteria were grown overnight at 37 ⁰ C. *E. coli* gDNA were extracted using TIANamp Bacteria DNA Kit. DpnI was bought from NEB and used as protocol described. All the samples and standards were loaded at 1 µg. 1% agarose gel electrophoresis was used to check digestion results.

### MeDIP-seq (6mA-IP-seq)

MeDIP-seq used in this paper was optimized from several protocols^13-15^. gDNA was extracted using TIANGEN DNASeure Plant Kit and then treated with RNase A overnight. Then the gDNA was diluted to 100 ng/µl with TE buffer, 100µl diluted gDNA was put in each tube and sonicated to 200-400 bp using Biorupter UCD-600. The 200-400 bp sized DNA was extracted using Takara Gel DNA Extraction Kit ver.4.0. DNA was denatured at 95 ⁰ C for 10 mins and chilled in ice immediately for 5 mins. 20 µl of denatured DNA was stored as input. The rest of the DNA was incubated with 3 µg 6mA antibody at 4 ⁰ C for more than 6 hours. Dyna beads (Thermofisher 10001D) were washed twice using 1×IP buffer and pre-blocked in 0.8 mL 1×IP buffer with 20 µg/µl BSA. Pre-blocked beads were washed twice using 1×IP buffer (5×IP buffer: 50mM Tris-HCl, 750mM NaCl and 0.5% vol/vol IGEPAL CA-630), and then incubated DNA-antibody was added to pre-blocked beads, and rotated overnight at 4 ⁰ C. The beads were washed 4 times for 10 mins with 1×IP buffer. IP products were suspended in 400 µL preheated elution buffer at 65 ⁰ C for 15 mins to yield the 6mA-IPed library; repeat this step using 300 µL preheated elution buffer (Elution buffer: 50 mM NaCl, 20 mM Tris-HCl, 5 mM EDTA, 1% SDS). Eluted DNA was combined and then added to an equal volume of phenol-chloroform-isopentanol, vortexed and centrifuged at 13000 rpm for 5 mins at room temperature. The aqueous phase was transferred into a new tube and mixed with an equal volume of ethanol to precipitate the eluted DNA. The library was prepared using VAHTS^TM^ Turbo DNA Library Prep Kit for Illumina and AHTS^TM^ Multiplex Oligos set 1 for Illumina. Sequencing was done by BGI (Shenzhen) and GENEWIZ (Suzhou).

### Phytophthora transformation

*Phytophthora* CRISPR/Cas9 gene editing and transformation was performed as previously described^50^. The sgRNA target sites were selected using an online tool (http://grna.ctegd.uga.edu/).

### RNA extraction, RNA-seq and qRT-PCR

Total RNA of 3-day-old *P. sojae* hyphae were isolated using Omega Total RNA Kit I according to the manufacturer’s manual. RNA quality was measured using Nanodrop ND-1000 and 1% agarose gel electrophoresis. RNA-seq service was provided by BGI and 1gene. RNA reverse transcription was conductd using Takara PrimerScript^TM^ RT reagent Kit with gDNA eraser. Quantitative RT-PCR was performed using the ABI PRISM 7500 Fast Real-Time PCR System.

### High-throughput sequence data analysis

RNA-seq data was mapped to *P. sojae* v1.1 using Tophat2, and MeDIP-seq data was mapped to *P. sojae* v1.1 using bowtie2. Gene expression data was generated by Cufflinks. MeDIP-seq data was normalized and visualized using deepTools^51^ and IGV^52^. 6mA methylation peaks were called using SICER. Figure of two-speed genome was produced ^as describe before^53^, 6mA distribution was calculated as log_2_(IP_RPKM_/input_RPKM_+1), RPKM^ value from the regions before TSS 500bp (real length and reads number will be calculated if the length of flanking intergenic regions<500bp). *Phytophthora* repeat sequences were referenced in previous publications^24^ and re-annotated here by RepeatMasker^54^.

## Acknowlegements

We thank Dahua Chen's Lab (Chinese Academy of Sciences) for 6mA blind test and shared 6mA quantification protocol. Jianzhao Liu (Zhejiang University), Sebastian Schornack (University of Cambridge) and Brett Tyler (Ohio State University) are also appreicated for helpful discussions. Ms. Hairong Xie (Nanjing Agricultural University) for MeDIP-seq, Ms Jiangyan Xu (Nanjing Agricultural University) for 5mC detection. Mark Gijzen (Agriculture and Agri-Food Canada), and The Sainsbury Laboratory student Aleksandra Bialas, Erin Zess and Jess Upson contributed to the editing of the manuscript. This work was supported by the Chinese National Science Fund (31422044), Chinese Thousand Talents Plan to S. D., and the Fundamental Research Funds for the Central Universities (KYTZ201403).

## Author contributions

H.C., L.Y.W., F.Z., X.L., S.O.O. and H.Y.M. performed experiments; H.D.S., H.C., F.M., W.W.Y. analyzed data; H.C., T.T.G., L.B.J., Y.F.W., S.K., Y.C.W. and S.M.D designed the experiments and discussed. S.M.D., H.C. and S.K. wrote the manuscript.

## Material & Correspondence

Dr. Suomeng Dong will take care of material request and correspondence. Address: Weigang NO.1, Xuanwu District, Nanjing, Jiangsu Province, China

